# Multiomic analysis of genes related to oil traits in legumes provide insights into lipid metabolism and oil richness in soybean

**DOI:** 10.1101/2024.05.02.592228

**Authors:** Dayana K. Turquetti-Moraes, Cláudio Benício Cardoso-Silva, Fabricio Almeida-Silva, Thiago M. Venancio

**Author notes:** TMV: Laboratório de Química e Função de Proteínas e Peptídeos, Centro de Biociências e Biotecnologia, Universidade Estadual do Norte Fluminense Darcy Ribeiro. Av. Alberto Lamego 2000, P5, sala 217, Campos dos Goytacazes, RJ, Brazil.

## Abstract

Soybean (*Glycine max*) and common bean (*Phaseolus vulgaris*) diverged approximately 19 million years ago. While these species share a whole-genome duplication (WGD), the *Glycine* lineage experienced a second, independent WGD. Despite the significance of these WGDs, their impact on gene families related to oil-traits remains poorly understood. Here, we report an in-depth investigation of oil-related gene families in soybean, common bean, and twenty-eight other legume species. We adopted a systematic approach that included transcriptome and co-expression analysis, identification of orthologous groups, gene duplication modes and evolutionary rates, and family expansions and contractions. We curated a list of oil candidate genes and found that 91.5% of the families containing these genes expanded in soybean in comparison to common bean. Notably, we observed an expansion of triacylglycerol (TAG) biosynthesis (∼3:1) and an erosion of TAG degradation (∼1.4:1) families in soybean in comparison to common bean. In addition, TAG degradation genes were two-fold more expressed in common bean than in soybean, suggesting that oil degradation is also important for the sharply contrasting seed oil contents in these species. We found 17 transcription factor hub genes that are likely regulators of lipid metabolism. Finally, we inferred expanded and contracted families and correlated these patterns with oil content found in different legume species. In summary, our results do not only shed light on the evolution of oil metabolism genes in soybean, but also present multifactorial evidence supporting the prioritization of candidates for crop improvement.

## 1. Introduction

The Fabaceae family (legumes) is a large group of flowering plants, with around 19,500 species. This family is remarkably diverse and stands out as the second most economically important family of crop plants after grasses (Azani et al. 2017; Thorne 2002). The notable grain legumes — such as chickpea (*Cicer arietinum*), pea (*Pisum sativum*), peanut (*Arachis hypogaea*), common bean (*Phaseolus vulgaris*), and soybean (*Glycine max*) — play a critical role in human and animal nutrition, as well as in industrial applications. For example, peanut and soybean are rich in oil and protein, while pea and common bean are rich in starch and protein (Aziziaram et al. 2021; Pattee et al. 1983; Shen, Hong, and Li 2022; Yao et al. 2020). Interestingly, oil content among legume species can vary dramatically, from ∼2% to ∼50% in common bean and peanut, respectively. Despite the differences in seed oil content, soybean and common bean are related crops that diverged approximately ∼19 million years ago (mya) (Lavin, Herendeen, and Wojciechowski 2005; Stefanović et al. 2009). These species shared a whole-genome duplication event (WGD, also referred to as polyploidization) ∼58 mya, while a second WGD (∼13 mya) occurred in the common ancestor of the *Glycine* genus, making soybean a suitable model for investigating the effects of WGD on gene family evolution (Shoemaker, Schlueter, and Doyle 2006).

Gene and genome duplications have been extensively associated with plant adaptation and diversification (Van de Peer, Mizrachi, and Marchal 2017; Zhuang et al. 2022). Gene duplication may occur through mechanisms such as WGDs or small-scale duplications (SSD) (Ren et al. 2018; Flagel and Wendel 2009). WGDs also played a role in the domestication of plants that eventually became modern crops, such as wheat and maize (Carretero-Paulet and Van de Peer 2020; Hake and Ross-Ibarra 2015; Rahman et al. 2020). WGDs are typically followed by genome fractionation and rearrangement, restoring bivalent chromosome pairing and disomic inheritance — a process also known as diploidization (Z. Li et al. 2021). Intriguingly, the prevalence of WGDs is significantly greater in plants than in other lineages of multicellular organisms (Panchy, Lehti-Shiu, and Shiu 2016). Gene duplicates that survive diploidization form families and often diverge at the sequence, epigenetic, and transcriptional levels (Freeling 2009), resulting in neofunctionalization or subfunctionalization. Neofunctionalization involves the acquisition of new functions, while subfunctionalization results in the division of the original function of the gene ancestor among its copies (Freeling 2009; Freeling, Scanlon, and Fowler 2015). Over time, gene families can gain or lose genes, generating a wealth of genetic material for adaptation (Cheng et al. 2018; Moharana and Venancio 2020; Renny-Byfield and Wendel 2014; Soltis et al. 2009). A comprehensive analysis of gene family evolution and expression data can contribute to the selection of candidates to improve oil-related traits. A remarkable example was the development of a lipoxygenase-free soybean, leading to improvements in the palatability of soybean oil and protein products. This advancement was achieved through the implementation of a pooled CRISPR-Cas9 system specifically targeting three soybean lipoxygenase genes from a set of 36 previously reported candidates (J. Wang et al. 2020; H. Song et al. 2016).

Although substantial efforts have been dedicated to the investigation of pivotal genes involved in oil content and quality (Borisjuk et al. 2005; B. Chen et al. 2020; Kanai et al. 2019; Li-Beisson et al. 2017; L. Lu et al. 2021; X. Lu et al. 2016; Manan et al. 2017; Marchive et al. 2014; Meinke, Chen, and Beachy 1981; Nguyen et al. 2016; Pham, Shannon, and Bilyeu 2012; Sandhu et al. 2007; Turquetti-Moraes et al. 2022), the evolution of these gene families and how WGDs shaped them remain largely unexplored. In the present study, we have undertaken an in-depth exploration of gene families associated with oil traits in legumes, with particular emphasis in soybean. We observed that the increase in soybean oil content was deeply impacted by the expansion of gene families shared with common bean. In addition, we hypothesize that most genes associated with lipid and fatty acid (FA) metabolism reverted to single copy after the ∼58 mya WGD and duplicated at the ∼13 mya WGD. In contrast, genes responsible for regulatory functions were often retained as duplicates in both species after the ∼58 mya WGD and duplicated again in the ∼13 mya WGD. Further, TAG degradation genes were two-fold more expressed in common bean than in soybean. Co-expression analysis uncovered 17 transcription factor (TF) hub genes that are strong candidate regulators of lipid metabolism. Finally, we inferred expanded and contracted orthologous groups and correlated these patterns with the oil contents found in different legume species. Our study expands the knowledge of several metabolic pathways, pinpoint key TFs, and show evidence for novel gene candidates involved in oil biosynthesis. Together, the results presented here also bear the potential to have practical applications by presenting the most promising targets to improve soybean oil content and quality according to the current landscape of genomics data.

## 2. Results and discussion

### 2.1 Soybean genes involved in oil traits belong to families shared with common bean

We selected 2,176 soybean genes related to oil traits as a reference to find candidate homologous gene families (Supplementary Table 1; see materials and methods). This set of genes, henceforth called oil genes, are distributed along 567 soybean homologous families, out of which 562 contain at least one common bean homolog (Supplementary Table 2). Only five families do not have a common bean homolog (HOM05D015518, HOM05D006604, HOM05D031525, HOM05D130031, and HOM05D039847) and include some genes (e.g. Glyma.02G006100, Glyma.02G281500, Glyma.08G064400, Glyma.16G133700, and Glyma.20G068000) reported as candidates to improve oil quality in soybean (Niu et al. 2020) (Supplementary Table 1; Supplementary Figure 1). These results indicate that the basic genetic machinery responsible for oil accumulation in soybean was already present in its last common ancestor shared with common bean, strongly suggesting that oil richness was acquired via expansions or contractions of extant gene families, mutations, and changes in transcriptional and epigenetic regulation, which were at least partially influenced by anthropogenic processes such as domestication and breeding (Turquetti-Moraes et al. 2022; J. Wang et al. 2020; M. Zhang et al. 2022).

About 14.57% (7,706 of 52,872) of the soybean genes and 15.44% (4,236 of 27,433) of the common bean genes belong to oil homologous families (Supplementary Table 3). These genes are enriched in functional terms related to metabolism and regulation of gene expression in both species (Supplementary Table 4), particularly TFs (e.g. GmDof4: Glyma.17G081800, GmDof11: Glyma.13G329000, GmMYB73: Glyma.06G303100, GmDREBL: Glyma.12G103100); response to oxidative stress (e.g. peroxidases: Glyma.19G066200, Glyma.07G263000); and metabolism of lipid and FAs (e.g. phospholipase: Glyma.03G159000, desaturase: Glyma.13G038600). Interestingly, the overexpression of GmMYB73 promotes lipid accumulation in soybean and its ectopic expression with other TFs (GmDof4, GmDof11, and GmDREBL) increased seed size/weight in transgenic *Arabidopsis* (Duan et al. 2023; Y.-F. Liu et al. 2014). In addition, 259 and 42 GO or Interpro terms were enriched only in soybean and common bean, respectively. For example, in soybean, we observed enrichment of terms including lipid glycosylation, FA and lipid metabolic process, and TAG biosynthetic process. On the other hand, in common bean, we found enrichment for lipid transport, microtubule nucleation and polymerization, and protein domains of lipoxygenases (Supplementary Table 4).

We investigated changes in sizes of the 562 gene families containing soybean and common bean genes (Figure 1). Using soybean as a reference, 2.1% (12), 91.5% (514), and 6.4% (36) of the families lost, gained, and had their sizes unchanged (i.e. neutral), respectively (Supplementary Table 2). Loss families were enriched in lipid metabolic process, especially to protein domains related to phospholipase/lysophospholipase. Neutral families were enriched in primary metabolic process, proteolysis, phospholipid biosynthetic processes, lipoate metabolic processes, lipid transport, and some enzyme domains or subunits (e.g. biotinyl protein ligase, FA desaturase, and seed storage helical domain) (Supplementary table 5). In gain families, by far the major group, we identified two scenarios: 25.5% (131 families) exhibited one common bean gene to two or more in soybean (1:2+), while 74.5% (383 families) had at least two common bean genes to three or more soybean homologs (2:3+). In summary, 1:2+ families were enriched in lipid and FA metabolism genes, while those families with duplications in common bean and new duplications in soybean (2:3+) were enriched in regulatory processes (e.g. gene expression and RNA metabolism) and response to stress (Supplementary Table 2; Supplementary Table 5). The 1:2+ families showed one peak in lower K_s_ values, while the 2:3+ families had two peaks. The peaks in both distributions correspond to the expected WGD K_s_ ages (Figure 1). Hence, we propose that most genes associated with lipid and FA metabolism (enriched in 1:2+ families) reverted to single copy after the ∼58 mya WGD and duplicated at the ∼13 mya WGD. In contrast, genes responsible for regulatory functions (enriched in 2:3+ families) were often retained as duplicates in both species after the ∼58 mya WGD and duplicated again in the ∼13 mya WGD. Although one cannot rule out the possibility of extreme sequence divergence and neofunctionalization, these results corroborate our hypothesis that the increase in oil content in soybean was deeply impacted by gene families shared with common bean that independently expanded in the *Glycine* WGD event.

**Figure 1.**
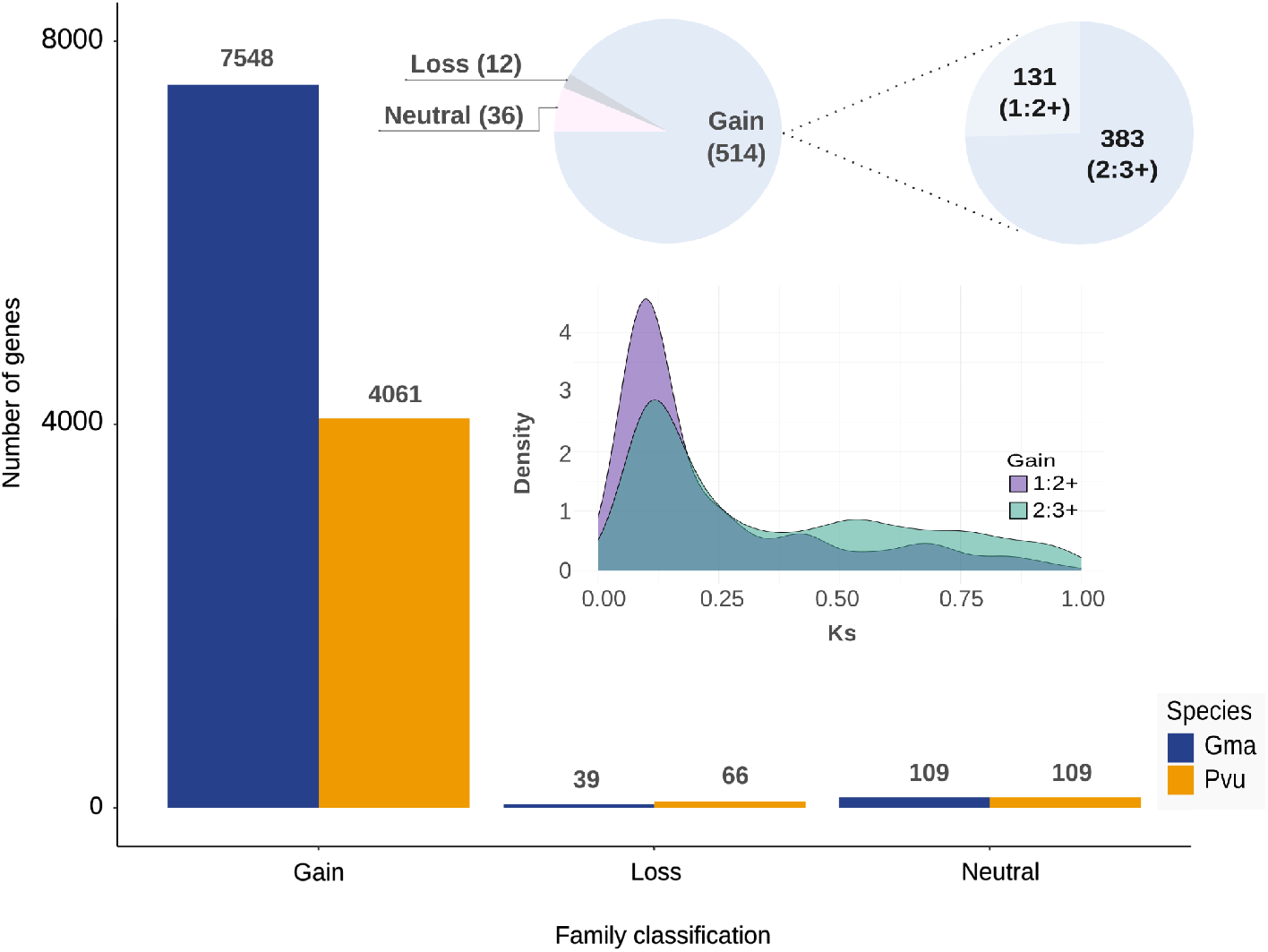
Number of soybean and common bean oil-related genes in different relative family size categories. Inset: number of families in each category. The two categories of gain families (1:2+ and 2:3+) are highlighted, alongside their respective K_s_ density plots. **Gain**: families with more genes in soybean (Gma) than in common bean (Pvu). **Loss**: families with less genes in Gma than in Pvu. **Neutral**: families with the same number of genes in both species.

### 2.2 Candidate oil families and expression patterns of genes from triacylglycerol (TAG) pathways

In plants, TAGs can be synthesized by two distinct routes. The classical Kennedy pathway involves the sequential acylation of glycerol-3-phosphate (G3P) (Figure 2.A). G3P is activated through acylation by acyl-CoA:glycerol-3-phosphate acyltransferase (GPAT), leading to the formation of lysophosphatidic acid (LPA); another acyl group is added to LPA by acyl-CoA:lysophosphatidic acid acyltransferase (LPAAT), forming phosphatidic acid (PA). PA is then dephosphorylated by PA phosphatase (PAP), leading to the formation of diacylglycerol (DAG). Finally, acyl-CoA:diacylglycerol acyltransferase (DGAT) adds the third acyl group to DAG, forming TAG. Alternatively, TAGs can be synthesized through the complex pathway, in which DAG originates from preexisting membrane lipids such as phosphatidylcholine (PC) through different pathways (Figure 2.A) (Bates 2016; Bates et al. 2009; Bates, Stymne, and Ohlrogge 2013). Over 90% of the acyl chains esterified to the glycerol backbone in developing soybean embryos originate from the complex pathway (Bates et al. 2009; Bates and Browse 2012). Hence, manipulating FA composition and seed oil content in soybean requires a deep understanding of these pathways. Furthermore, given its paleopolyploid genome, pinpointing the paralogs that are truly involved in seed lipid metabolism imposes an additional layer of complexity.

**Figure 2.**
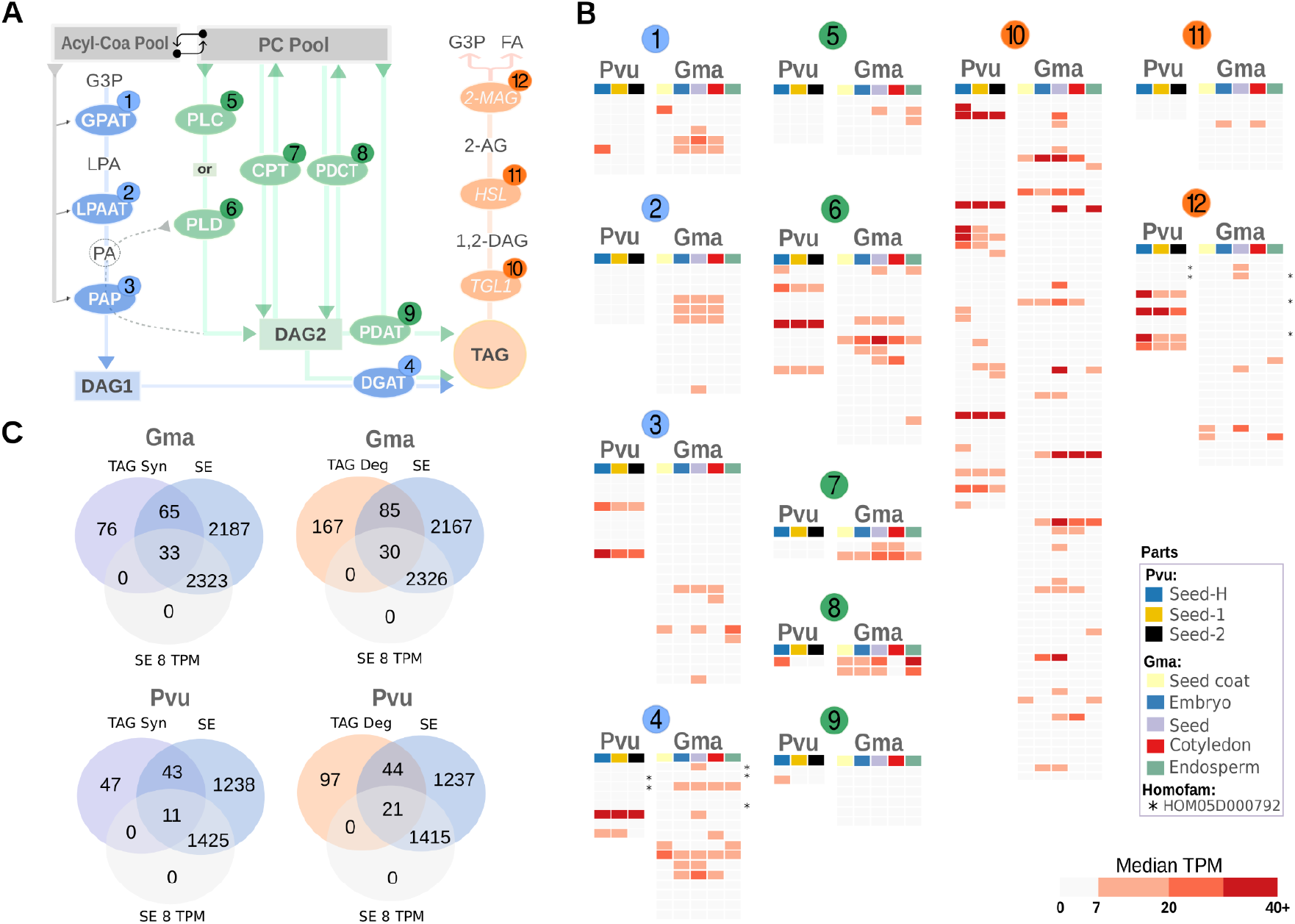
Major enzymes involved in DAG/TAG synthesis and degradation. **A**. Reactions involved in TAG degradation and formation from FAs. Blue: Kennedy pathway (gray arrows indicate the source of FA to feed the pathway). Green: complex pathway; the dashed arrow indicates a reaction PLD-PAP to form DAG from PC; the arrows from CPT and PDCT indicate a reversible reaction for the formation of PC-derived DAG. Orange: TAG degradation pathway. **B**. Expression of soybean and common bean genes encoding enzymes in each step of the pathways from panel A. Asterisks (*) denote unannotated genes from the HOM05D000792 family, which comprises genes associated with TAG synthesis and degradation pathways. **C**. Venn diagram of gene counts in TAG homologous families and their seed expression in Gma and Pvu. Abbreviations: **G3P**: glycerol-3-phosphate; **GPAT**: acyl-CoA glycerol-3-phosphate acyltransferase; **LPA**: lysophosphatidic acid; **LPAAT**: acyl-CoA lysophosphatidic acid acyltransferase; **PA**: phosphatidic acid; **PAP**: phosphatidic acid phosphatase; **DAG**: diacylglycerol; **DAG1**: DAG from the Kennedy pathway; **DAG2**: PC-derived DAG; **DGAT**: acyl-CoA diacylglycerol acyltransferase; **TAG**, triacylglycerol; **PC Pool**: phosphatidylcholine pool; **PLC**: phospholipase C; **PLD**: phospholipase D; **CPT**: cytidine diphosphate-choline diacylglycerol cholinephosphotransferase; **PDCT**: phosphatidylcholine diacylglycerol cholinephosphotransferase; **PDAT**: phospholipid diacylglycerol acyltransferase; **TGL1**: triacylglycerol lipase; **HSL**: hormone-sensitive lipase; **2-MAG**: 2-monoacylglycerol acylhydrolase; **FA**: fatty acid; **Pvu**: common bean; **Gma**: soybean; **TPM**: transcripts per million; **Seed-H**: heart stage seeds (∼7 mg); **Seed-1**: state 1 seeds (∼50 mg) ; **Seed-2**: stage 2 seeds (∼150 mg). **TAG Syn**: TAG synthesis pathway; **TAG Deg**: TAG degradation pathway; **SE**: seed expression (>=1 TPM); **SE 8 TPM**: seed expression (>=8 TPM). The genes in this heatmap can be found in Supplementary Table 7 and Supplementary Table 8. Venn diagram was generated using Venn diagram (https://bioinformatics.psb.ugent.be/webtools/Venn/).

By meticulously curating homologous families from SoyCyc TAG pathway genes (soybase.org), we found 352 TAG genes in soybean (Supplementary Table 6), which are distributed in 43 families. These families include 204 and 116 genes with seed expression (TPM > 1) in soybean and common bean, respectively (Figure 2.B, Supplementary Table 7, Supplementary Table 8). We found one family, HOM05D000792, comprising genes belonging to TAG synthesis (e.g. DGAT: Glyma.16G051200, Glyma.16G051300) and degradation (e.g. 2-MAG: Glyma.03G243700, Glyma.19G241200). Other five HOM05D000792 members that are expressed in seeds encode WD repeat-containing proteins with unknown functions (e.g. Glyma.19G241800, Glyma.03G244500, Glyma.10G159000, Phvul.006G097800, and Phvul.006G098200). We kept these five genes in both TAG biosynthesis and degradation pathways (asterisk marks in Figure 2.B, Supplementary Table 7 and Supplementary Table 8).

The families encoding enzymes from each step of the TAG pathways presented at least one gene with expression of ∼10 TPM in soybean seeds, except for phospholipid diacylglycerol acyltransferase (PDAT) (Figure 2.B). Thus, in order to select the most promising candidate genes, we adopted a threshold of 8 TPM (Supplementary Figure 2), which allowed us to find 33 and 11 (∼3:1) TAG synthesis and 30 and 21 (∼1.4:1) TAG degradation genes expressed in soybean and common bean seeds, respectively (Figure 2.C, Supplementary Table 7 and Supplementary Table 8). These results suggest both a relative expansion in the TAG biosynthesis and an erosion of TAG degradation components in soybean when compared to common bean, even if we consider the 2:1 ratio expected because of the ∼13 mya WGD. We also analyzed the expression of these genes in all plant parts (Supplementary Figure 3, Supplementary Figure 4). Four GmGPATs and only one PvGPAT had at least 8 TPM in seeds (Glyma.07G069700, Glyma.05G131100, Glyma.08G085800, Glyma.09G119200, Phvul.010G099700) (Figure 2.B). Liu et al. showed that only GmGPAT9-2 (Glyma.09G119200) out of sixteen tested GmGPATs with high acyltransferase activity may not play a direct role in TAG formation. However, they found that seed-specific expression of GmGPAT9-2 in *Arabidopsis* increased the proportion of arachidic acid (C20:0) and erucic acid (C22:1) without an increase in the total oil content (H. Liu et al. 2022). Except for Glyma.08G085800, the GmGPATs mentioned above were corroborated by Liu et al. However, we found that this gene is highly expressed in seeds, specially in embryo and cotyledons (Figure 2.B, Supplementary Table 7), supporting its role in TAG synthesis.

In a second group of transferases, LPAATs regulate the synthesis of PA, an intermediate in the formation of membrane, signaling, and storage lipids (Kim and Wang 2020). From the four GmLPAATs reported here (Figure 2.B, Supplementary Table 7), Glyma.10G095500 was highly expressed in cotyledons, corroborating previous studies (X. Wang et al. 2019). Glyma.02G181300 and Glyma.12G163500 are within QTL associated to phosphatidylcholine (qPC-2.1) and phosphatidylinositol (qPI-12.1), respectively (Anshu et al. 2022) (Supplementary Table 7). Finally, DGAT forms an ester linkage between a fatty acyl-CoA and the DAG free hydroxyl group (G. Chen et al. 2022). From a set of 26 recently studied GmDGATs (S. Zhao et al. 2023), we found six showing high expression in seeds (Glyma.16G051200, Glyma.01G156000, Glyma.16G115700, Glyma.13G118300, Glyma.09G065300, and Glyma.13G106100). All these GmDGATs appear to be involved in TAG assembly, although DGAT1s such as GmDGAT1A (Glyma.13G106100) and GmDGAT1C (Glyma.09G065300) influence oil content and quality more prominently (Torabi et al. 2021; J. Zhao et al. 2019).

The complex pathway for DAG/TAG synthesis (Figure 2.A) is influenced by phospholipase C (PLC), phospholipase D (PLD) associated with PAPs (PA phosphatase), PDAT, and CPT or phosphatidylcholine diacylglycerol cholinephosphotransferase (PDCT). PLC hydrolyzes PC to DAG and phosphocholine, while PLD hydrolyses the choline head-group of PC and forms PA, an intermediate to the synthesis of other phospholipids. Subsequently, PAP removes the phosphate head-group of PA, converting it to DAG (Figure 2.A) (Bates, Stymne, and Ohlrogge 2013; Lee et al. 2011; W. Yang et al. 2017; Wakelam 1998). In soybean seeds, PLD is generally more expressed than PLC (Figure 2.B), although the PLCs Glyma.04G196700 and Glyma.06G169100 may be relevant candidates due to their endosperm expression (Figure 2.B). From the six GmPLD genes analyzed here (Glyma.01G162100, Glyma.07G031100, Glyma.08G211700, Glyma.11G081500, Glyma.13G364900, Glyma.15G267200), PLDα1 (Glyma.13G364900) is a promising target considering its high expression in cotyledons (∼20 TPM). Further, suppression of PLDα in soybean results in decreased levels of polyunsaturated FAs in TAG (Lee et al. 2011), suggesting a flow from PC to TAG via PLD-PAP without PDCT. In addition, GmPLDα3 is also highly expressed in seeds (Glyma.08G211700: cotyledon expression of ∼28 TPM) and was reported as associated with malate (J.-Y. Liu, Li, et al. 2020). Considering the expression of GmPAPs (Figure 2, Supplementary Figure 3) we suggest Glyma.10G270000 as a promising candidate to integrate this route.

Other potential routes for DAG2/TAG formation involves the reverse activity (i.e. PC to DAG) of PDCT or amino alcohol phosphotransferase (AAPT) also known as cytidine diphosphate-choline:diacylglycerol cholinephosphotransferase (CPT). These enzymes synthesize unique DAG and PC molecular species. PDCT catalyzes the interconversion between PC and DAG, contributing to the enrichment of polyunsaturated FAs in TAGs. The conversion of PC to DAG varies even between closely related species. PDCT transfers ∼40% of oleic acid from PC to DAG in *Arabidopsis* against ∼18.2% in canola (S. Bai et al. 2020; C. Lu et al. 2009). We propose the GmPDCTs Glyma.07G029800 and Glyma.08G213100 as interesting candidates because of their high expression in seeds (Supplementary Table 7). CPTs GmAAPT1 (Glyma.02G128300) and GmAAPT2 (Glyma.12G081900) were recently reported as crucial enzymes in TAG metabolism (Y. Bai et al. 2021), which is in line with the high GmAAPT2 expression in seeds. Finally, in addition to DGAT, PDAT is also involved in TAG assembly. GmPDAT (Glyma.13G108100) catalyzes the transfer of a FA moiety from the sn-2 position of a phospholipid to the sn-3-position of sn-1,2-DAG, forming TAG and a lysophospholipid (Pan, Peng, and Weselake 2015). PDATs often have contrasting expression profiles in different plant species (Torabi et al. 2021; Pan, Peng, and Weselake 2015). For example, PDAT was more expressed in plants that accumulate epoxy and hydroxy FAs (e.g. *Vernonia galamensis* and *Erysimum lagascae*) than in soybean and *Arabidopsis* (R. Li, Yu, and Hildebrand 2010). GmPDAT has been reported as associated with acyl-lipid metabolism and likely interacts with GmDGAT1 (Xu et al. 2018), although further studies are warranted to better understand this interaction and its roles in TAG synthesis (J.-Y. Liu, Zhang, et al. 2020).

TAG degradation during seed development is also important for the oil content of mature seeds (Ding et al. 2019). Strikingly, TAG degradation genes exhibited significantly higher expression levels in common bean (mean ∼47 TPM) than in soybean (mean ∼19 TPM). Furthermore, these genes exhibit high expression levels even in the early stages of seed development. We have identified TAG degradation genes that could be tested in soybean, out of which we highlight GmTGL1s (Glyma.01G067200, Glyma.02G043300, Glyma.06G294900, Glyma.04G255500, and Glyma.09G233900) (Supplementary Figure 3). In common bean, certain TGL1 genes, such as Phvul.001G168200, Phvul.011G003600, Phvul.011G088700, Phvul.005G114300, and Phvul.010G098300 were highly expressed in seeds (Supplementary Figure 4). Notably, Phvul.011G003600 demonstrated consistently high expression across all seed stages. These findings suggest that the high expression of TAG degradation genes is a key factor to the lower oil accumulation in common bean. In conclusion, achieving high TAG levels in seeds requires an intricate system involving reduced TAG degradation, increased de novo FA synthesis, increased DAG production, and a flow from PC to combine saturated and unsaturated chains. Investigating the concerted action of these genes is key for enhancing oil content and quality in soybean seeds.

### 2.3 Identification of oil candidate genes with high expression in seeds

In addition to the detailed TAG pathways reported above, we applied the same threshold of 8 TPM in seeds to mine new genes potentially involved in oil metabolism. Approximately 50.7% (17,086/33,684) and 50.2% (8,941/17,805) of the genes expressed in seeds or seed subregions (i.e., endosperm, cotyledon, embryo or seed coat) showed at least 8 TPM in soybean and common bean, respectively (Supplementary Table 9, Supplementary Table 10). Expectedly, these genes are enriched in the metabolism of various molecules such as nitrogen, peptide, amino acid, ribose phosphate, nucleotide, ATP, as well as in translational elongation, protein transport, and gene expression. Out of the 17,086 genes highly expressed in soybean seeds, 562 were seed-specific and are enriched in defense responses and negative regulation of protein metabolic processes (e.g. proteolysis and peptidase activity), essential during seed development and other processes (Santamaría et al. 2014) (Supplementary Table 11). Considering only genes from candidate oil homologous families, we found 1.64 times more soybean (2,356) than common bean (1,436) genes expressed in seeds (Supplementary Table 9, Supplementary Table 10). These genes are distributed in 510 homologous families that can be classified in three categories: I. 402 families with at least one member expressed in soybean and common bean; II. 90 families with members expressed in soybean but not in common bean and; III. 18 families with members expressed in common bean but not in soybean (Supplementary Table 12).

The 402 homologous families from category I comprise 2,198 and 1,413 genes expressed in soybean and common bean, respectively. These families are enriched in a myriad of functions (Supplementary Table 11) and also encompass strong candidates for oil accumulation, such as GmSEIPIN1A (Glyma.09G250400), involved in TAG accumulation and lipid droplet assembly, maintenance, and proliferation (Pyc et al. 2021; Qi et al. 2023; Taurino et al. 2018).

The second category comprises 90 families containing genes expressed in soybean but not in common bean seeds. These genes emerge as promising candidates to account for the contrasting lipid contents found in soybean and common bean. Interestingly, these families are enriched in various lipid metabolism processes (e.g. phospholipid/glycerol acyltransferase and CDP-alcohol phosphatidyltransferase) (Supplementary Table 11). Although most of these genes are expressed in different plant parts, 24 are predominantly expressed in seeds and seed subregions, including lipases (Glyma.03G256900 and Glyma.19G255100), Zinc-finger-CW domain proteins (Glyma.14G204400 and Glyma.18G052100), and a sterol dehydrogenase (Glyma.06G058200) (Supplementary Figure 5). In addition, these families also comprise poorly characterized genes (HOM05D015518: Glyma.20G068000, HOM05D025622: Glyma.05G011200, HOM05D031525: Glyma.08G064400, HOM05D039847: Glyma.02G281500, HOM05D130031: Glyma.02G006100) reported as candidates to improve oil content and quality (Niu et al. 2020; Fang et al. 2017) (Supplementary Figure 5). Together, these results support the recruitment of multiple genes to soybean seed metabolism after the split between soybean and common bean.

The third category comprises 18 homologous families with genes expressed in common bean but not in soybean seeds (Supplementary Figure 6). For example, the TFs Phvul.007G171333, Phvul.006G179900, and Phvul.009G149200 are highly expressed in the heart seed stage. The phosphatidylinositol PLC (Phvul.009G025300), associated with plant immunity (Tasma et al. 2008), also appears to play roles throughout seed development. In general, genes from this category suggest a deviation from TAG storage (Supplementary Figure 6). For example, Phvul.002G054200 encodes an enzyme with domains associated with wax ester synthesis and acyl group transfer, supporting a role in lipid modification instead of FA storage. These genes may drive FA metabolism towards structural lipid modification or secondary metabolism rather than storage.

About 3.5% (83) of the soybean oil candidate genes showed seed specific expression (Supplementary Table 9). In order to have a more conservative estimate, we retrieved the tissue specificity metrics from a broader and more heterogeneous set of samples available at the Soybean Expression Atlas (Almeida-Silva, Pedrosa-Silva, and Venancio 2023). These 83 genes were enriched in protein domains such as GDSL lipase/esterase and MADS-box TFs (Figure 3, Supplementary Table 11). GDSL-type esterase/lipase proteins belong to the SGNH hydrolase superfamily that can hydrolyze various substrates, including thioesters, acyl esters, phospholipids, and amino acids (Akoh et al. 2004; Su et al. 2020). Out of the 194 previously identified GmGDSL-type genes (Su et al. 2020), eight were found here as seed-specific and are strong candidates to improve oil content (Figure 3). MADS-box TFs are widely known for their roles in flowering, growth, and development (Shu et al. 2013; Zeng et al. 2018). Thirteen GmMADS-box TFs were preferentially expressed in seeds (Figure 3). Studies in *Arabidopsis* and oil palm (*Elaeis guineensis Jacq.*) support MADS-box TFs as important regulators of lipid metabolism and responsible for a decreased accumulation of polyunsaturated fatty acids (S.-Y. Li et al. 2020; Sun et al. 2020). GDSL and MADS-box genes account for 25.3% of the soybean seed-specific genes with at least 8 TPM. Interestingly, the MADS-box Glyma.04G257100 clusters with the sucrose/hexose transporter GmSWEET24/GmSWEET10b (Glyma.08G183500) (Figure 3), known to influence the distribution of sugars from the seed coat to the embryo and playing a crucial role in key soybean seed traits, such as size, oil and protein contents (S. Wang et al. 2020; S. Lu et al. 2022). We also found seed-specific expression of GmSWEET23 (Glyma.08G077200), GmSWEET4 (Glyma.04G198400), and GmSWEET39/GmSWEET10a (Glyma.15G049200), which were also associated with seed oil and protein content (Hooker et al. 2022; C. Liu et al. 2023; H. Yang et al. 2019; Boyang, Wenlong, and Caiying 2023). Finally, we highlight the seed-specific expression of the gibberellin 20-oxidases (GmGA20OX) Glyma.03G019800 and Glyma.07G081700 (Figure 3). GmGA20OX overexpression in *A. thaliana* has been shown to increase seed weight and oil (X. Lu et al. 2016). In this context, it is important to mention that seed size and weight have been related to oil content and several genes associated with these traits have been identified in soybean (Alam et al. 2022; J. Li et al. 2019).

**Figure 3.**
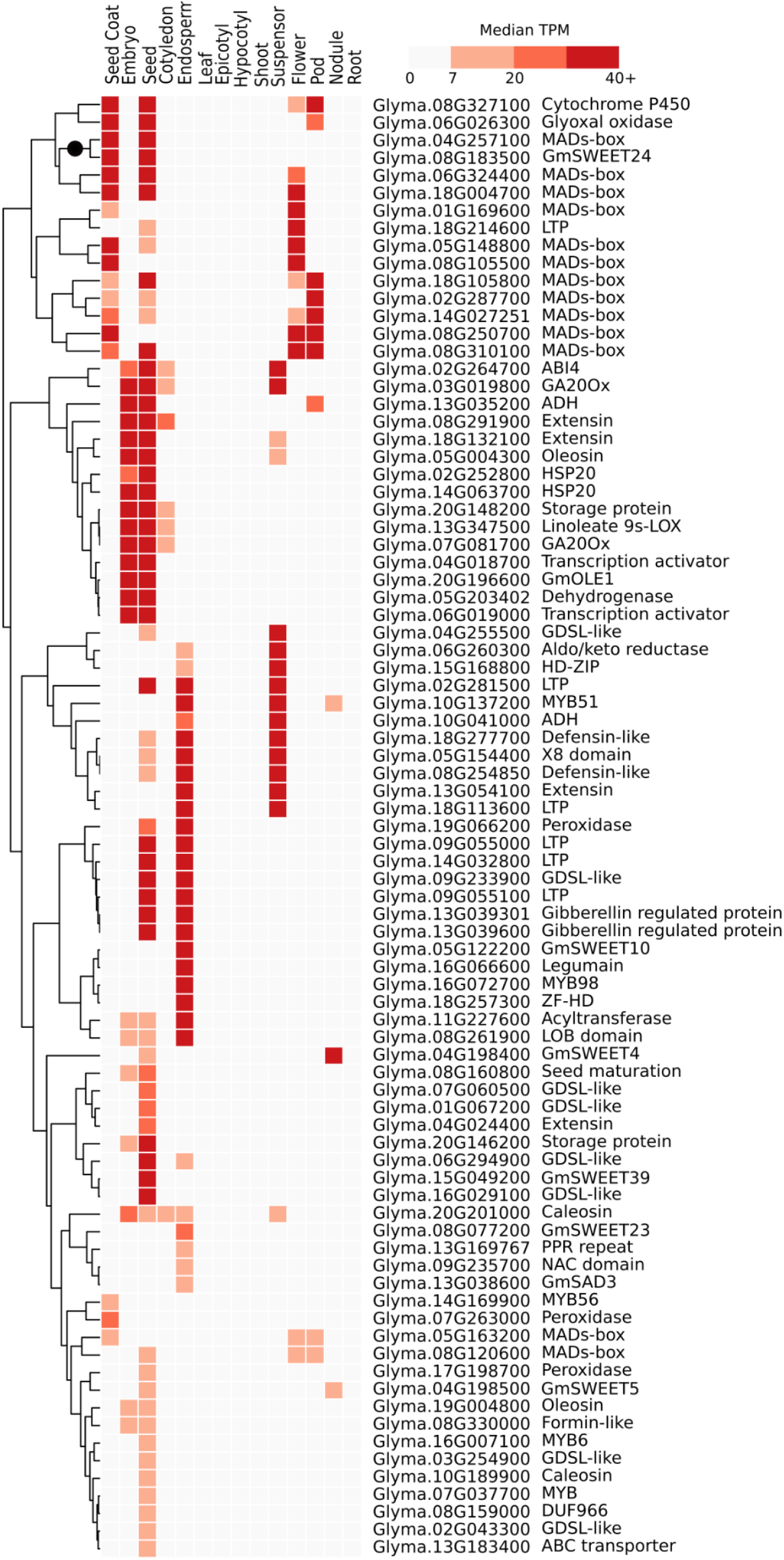
Expression of oil candidate genes with preferential expression in seeds. Tissue specificity metrics were obtained from the Soybean Expression Atlas (Almeida-Silva, Pedrosa-Silva, and Venancio 2023). Annotations were retrieved from Phytozome (V13) and Soybase.org. TPM: transcripts per million. Black circle marks the cluster with the GmSWEET24 and MADS-box genes discussed in the text.

### 2.4 Co-expression network of soybean genes with high seed expression from gain families

Aiming to better characterize novel seed oil genes, we used BioNERO (Almeida-Silva and Venancio 2022)(see methods for details) to compute a co-expression network (Almeida-Silva et al. 2020; Schaefer, Michno, and Myers 2017) of the gain-family genes with at least 8 TPM in seeds. Out of the 2,269 input genes, 92.9% (2,107) were distributed across 12 co-expression modules (Supplementary Figure 7). Except for the green module, we observed functional enrichment in all other modules: signal transduction and molecular transport, such as lipid transport (blue); cellular process, particularly photosynthesis and oil body stabilization (grey); cellular stress responses (greenyellow, brown); gene expression regulation (purple, magenta); multicellular organism development, cellular signaling and cytoskeletal dynamics. (tan, light green, dark red); protein processes/activity (light yellow); and lipid and FA metabolism (cyan); (Supplementary Table 13).

The cyan module comprises 33 TFs (Supplementary table 14) and important genes involved in seed oil quality and accumulation, such as GmFATA1A (Glyma.18G167300), GmFATA1B (Glyma.08G349200), GmDGAT1C (Glyma.09G065300), biotin carboxyl carrier protein (Glyma.18G243500, Glyma.09G248900), biotin carboxylase (Glyma.08G027600, Glyma.05G221100), ketoacyl-ACP synthase (KASI: Glyma.08G084300; KASII: Glyma.17G047000; KAS III: Glyma.09G277400), and Long-chain Acyl-Coa Synthetase (LACS: Glyma.06G112900, Glyma.13G079900, Glyma.20G060100) (Torabi et al. 2021; X. Wang et al. 2019). Glyma.07G110900 and Glyma.06G122600 encode a cytochrome P450 and an alcohol dehydrogenase, respectively. A previous study suggested that relative expression of these two genes promote the synthesis of linolenic acid in mature soybean seeds (X. Wang et al. 2019). In addition, we encountered four unannotated genes (Glyma.12G105300, Glyma.10G277900, Glyma.04G044200, and Glyma.05G141600) with potential activity in lipid metabolism (Supplementary Table 14).

Aiming to identify genes that may play significant roles in oil regulation and synthesis we analyzed modules that exhibit closest co-expression with the cyan module, i.e., darkred, green, and magenta (Supplementary Figure 8). We identified 15, 6 and 19 TF families containing 63, 8 and 52 genes distributed in darkred, green and magenta modules, respectively (Supplementary Table 14). Approximately 25% (31) of them (Glyma.02G016100, Glyma.02G274600, Glyma.02G303800, Glyma.04G010300, Glyma.04G044900, Glyma.04G050300, Glyma.05G098200, Glyma.05G140400, Glyma.05G175600, Glyma.06G079800, Glyma.08G132800, Glyma.08G360200, Glyma.09G241800, Glyma.10G016500, Glyma.11G242200, Glyma.12G040600, Glyma.13G153200, Glyma.13G202300, Glyma.14G041500, Glyma.14G071400, Glyma.14G205600, Glyma.16G011200, Glyma.16G012600, Glyma.16G152700, Glyma.17G096700, Glyma.17G132600, Glyma.17G157600, Glyma.17G174900, Glyma.18G014900, Glyma.19G022200, Glyma.20G200500) were previously reported as candidates for oil accumulation (Niu et al. 2020). Interestingly, within the magenta module, we found two NF-Y TFs (Glyma.12G236800 and Glyma.13G202300) as potential regulators of seed traits. Although NF-Y TFs are important for oil biosynthesis in *E. guineensis* (Yeap et al. 2017), their roles in soybean lipid metabolism remain unclear. Understanding the effects of gene silencing on TF regulation is also important. For example, we found five TFs (Glyma.01G081100, Glyma.04G010300, Glyma.14G112400, Glyma.17G096700; in darkred and Glyma.16G179900; in magenta) potentially regulated by DNA methylation during seed maturation (An et al. 2017).

We identified 189 hubs (Supplementary Table 14), of which 3.17% (6) are involved in TAG pathways (GmGPAT: Glyma.07G069700, GmPLD: Glyma.07G031100, PLDα1: Glyma.13G364900, and TGL1: Glyma.14G050600, Glyma.20G127800). The WD-repeat Glyma.03G244500 discussed in section 2.2 was also found as a hub and showed a remarkable expression correlation (∼0.8) with DGAT2A (Glyma.01G156000) (Supplementary Table 15). A recent study showed that DGAT2A enhances oil and linoleic acid contents in soybean seeds (Jing et al. 2021), highlighting the potential involvement of Glyma.03G244500 in TAG synthesis. In addition, we found three hubs encoding proteins of unknown function in the modules blue (Glyma.11G222000), green (Glyma.10G222300), and greenyellow (Glyma.13G092300), all connected to TFs and lipid metabolism genes (Supplementary Table 15). Interestingly, Glyma.13G092300 was previously reported as a candidate to improve oil quality (Niu et al. 2020). This gene is connected with MOTHER-OF-FT-AND-TFL1 (GmMFT: Glyma.05G244100; Supplementary Figure 9), which was proposed as a major gene of the classical QTL qOil-5-1 that regulates seed oil and protein content (Fang et al. 2017; J. Huang et al. 2020; Zhou et al. 2015; Cai et al. 2023). The connections of these three hubs with relevant genes to lipid metabolism and TFs support their importance in oil-related traits. Finally, we found 17 TF hub genes belonging to ten families, which might constitute bona-fide regulators of transcriptional programs involved in oil accumulation (Supplementary Table 14).

### 2.5 Conservation of oil candidate genes in legumes

We employed Orthofinder (Emms and Kelly 2019) to investigate the conservation of candidate genes across 30 legume species (Table 2, Supplementary Table 16). Approximately 94.7% (1,086,739/1,147,876) of the legume genes are distributed in 46,003 orthologous groups (OGs). Around 5.5% (63,594) of the genes are distributed in 12,249 species-specific OGs (Supplementary Table 16, Supplementary Table 17). *M. truncatula*, *P. sativum*, *L. japonicus*, *A. hypogaea*, and *S. tora* had the greatest frequencies of species specific OGs, while *A. hypogaea*, *A. ipaensis*, and *L. angustifolius* exhibited the highest frequencies of species-specific duplications (Supplementary Figure 10).

**Table 2.** Plant species used in this study, genome information and data source.

| Species | Assembly | Genome size | Publication | Database |
| --- | --- | --- | --- | --- |
| *Arachis duranensis* | V2.0 | 1.25 Gb | (Garg et al. 2022) | Legumepedia |
| *Arachis hypogaea* | V1.0 | 2.7 Gb | (Bertioli et al. 2016) | Phytozome 13 |
| *Arachis ipaensis* | V1.0 | 1.56 Gb | (Q. Lu et al. 2018) | DRYAD |
| *Cajanus cajan* | V2.0 | 833 Mb | (Garg et al. 2022) | Legumepedia |
| *Cercis canadensis* | V1 | 342 Mb | (Griesmann et al. 2018) | GigaDB |
| *Chamaecrista fasciculata* | V1 | 429 Mb | (Griesmann et al. 2018) | GigaDB |
| *Cicer arietinum* | V2.0 | 738 Mb | (Garg et al. 2022) | Legumepedia |
| *Cicer reticulatum* | — | 416 Mb | (S. Gupta et al. 2017) | NCBI |
| *Faidherbia albida* | — | 654 Mb | (Chang et al. 2019) | GigaDB |
| *Glycine max* | V4.0 | 978 Mb | (Schmutz et al. 2010) | Phytozome 13 |
| *Glycine soja* | V1.1 | 985 Mb | (Valliyodan et al. 2019) | Phytozome 13 |
| *Lablab purpureus* | — | 615 Mb | (Chang et al. 2019) | GigaDB |
| *Lotus japonicus* | V3.0 | 472 Mb | (Sato et al. 2008) | Kazusa |
| *Lupinus albus* | V1 | 450 Mb | (Hufnagel et al. 2020) | Phytozome 13 |
| *Lupinus angustifolius* | V1.0 | 924 Mb | (Hane et al. 2017) | NCBI |
| *Medicago truncatula* | V4.0 | 411 Mb | (Tang et al. 2014) | Phytozome 13 |
| *Mimosa pudica* | V1 | 557 Mb | (Griesmann et al. 2018) | GigaDB |
| *Nissolia schottii* | V1 | 466 Mb | (Griesmann et al. 2018) | GigaDB |
| *Phaseolus acutifolius* | V1.0 | 512 Mb | (Moghaddam et al. 2021) | Phytozome 13 |
| *Phaseolus lunatus* | V1 | 546 Mb | — | Phytozome 13 |
| *Phaseolus vulgaris* | V2.0 | 537 Mb | — | Phytozome 13 |
| *Pisum sativum* | V1 | 4 Gb | (Kreplak et al. 2019) | URGI/INRA |
| *Senna tora* | V1 | 547 Mb | (S.-H. Kang et al. 2020) | NCBI |
| *Spatholobus suberectus* | V1 | 793 Mb | (Qin et al. 2019) | NCBI |
| *Trifolium pratense* | V2 | 345 Mb | (De Vega et al. 2015) | Phytozome 13 |
| *Trifolium subterraneum* | V2.0 | 540 Mb | (Garg et al. 2022) | Legumepedia |
| *Vigna angularis* | V1.1 | 466 Mb | (K. Yang et al. 2015) | NCBI |
| *Vigna radiata* | V1 | 463 Mb | (Y. J. Kang et al. 2014) | NCBI |
| *Vigna subterranea* | — | 535 Mb | (Chang et al. 2019) | GigaDB |
| *Vigna unguiculata* | V1.2 | 519 Mb | (Lonardi et al. 2019) | Phytozome 13 |

We used the 2,269 soybean gain-family genes with at least 8 TPM in seeds as references to find 1,104 candidate OGs containing 67,577 genes. In general, approximately 47.7% (527) of the OGs are shared by all species (Supplementary Table 18). Expectedly, these core legume genes are enriched in essential metabolic pathways such as phosphatidate metabolism, glycolysis, tricarboxylic citric acid (TCA) cycle, ureide biosynthesis, and other energy metabolism pathways. The remaining 52.3% (577) OGs are enriched in stress response signaling, antioxidant defense mechanisms, and synthesis of bioactive compounds with potential medicinal applications, such as divinyl ether, chlorogenic acid, and justicidin (Grechkin 2002; A. Gupta et al. 2022; Hemmati and Seradj 2016) (Supplementary Table 19).

Three OGs (OG0045944, OG0035121 and OG0018534) contained genes only from two species known for their high oil content, namely soybean and peanut. The genes in these OGs are annotated as amidases (OG0045944: Glyma.08G054300, Glyma.08G197900); and auxin response factor - ARF (OG003512: Glyma.07G134900, AVLU3S; and OG0018534: Glyma.12G153700, AQQ3N0, FQ0W8J, K35AK0, MJ6K6I, NZ22HG, RIMX1M, RZFM32, Y8ANLA, YS88LH, YTF73Z). Interestingly, auxin can alter FA content and composition in soybean and microalgae (W. Liu, Hildebrand, and Collins 1995; Jusoh et al. 2015). Thus, these ARF genes into OG003512 and OG0018534 may be candidates to influence seed development and oil content. We also investigated the 1,104 OGs mentioned above for the presence of TAG pathway genes. We found 2,397 genes distributed across 41 OGs, out of which 19 are shared by all species (Supplementary Table 18). Among OGs not shared by all species, OG0027103 and OG0018208 contain fewer than 10 species. OG0027103, with HSL genes, is specific to *G.soja* and *G.max*; while OG0018208, with PLD genes, is specific to *A. hypogaea*, *C. fasciculata*, *G. soja*, *L. japonicus*, *M. truncatula*, *N. schottii*, *P. sativum* and *G. max*.

We found 46 OGs containing genes with seed-specific expression in soybean, of which 25 are shared by all species (Supplementary Table 20) and associated with regulatory processes (e.g.: metabolic, biosynthetic, and transcriptional), while the remaining 21 are enriched in lipoxygenases, divinyl ether biosynthesis, oleosin, SWEET transporters, among others (Supplementary Table 21). Some studies indicate that lipoxygenases in mature seeds produce conjugated unsaturated FA hydroperoxides, resulting in volatile compounds linked to the undesirable beany flavor (Rackis, Sessa, and Honig 1979; J. Wang et al. 2020). Other studies suggest that SWEET transporters significantly influence seed oil and protein contents (Duan et al. 2023; S. Wang et al. 2020). These results provide some genes that are likely involved in FA oxidation and nutrient uptake, which are important for seed quality and nutritional diversity found in legumes.

Finally, we employed CAFE5 (Mendes et al. 2020), a method based on gene birth (*λ*) and death (μ) rates, to investigate size changes in OGs. We ran CAFE only with the 1,104 OGs containing at least one of the 2,269 gain-family genes with a minimum expression of 8 TPM in seeds. From these OGs, 163 had significant contractions or expansions (p-value < 0.05, Figure 6). Out of these, 105 are shared by all species (Supplementary Table 18) and contain soybean genes related to lipid metabolism. Notable examples of these OGs include: OG0000430 (NADP-malic enzyme - NADP-ME: Glyma.13G354900; Glyma.15G019300), OG0000544 (GmbZIP123: Glyma.06G010200), OG0000840 (GmSWEET: Glyma.08G183500; Glyma.15G049200), OG0002123 (GmDGAT: Glyma.09G065300; Glyma.13G106100), and OG0000341 (containing the MADS box Glyma.04G257100, indicated here as a promising regulator of FA metabolism) (Torabi et al. 2021; S. Wang et al. 2020; Morley et al. 2023; Q.-X. Song et al. 2013). Conversely, among the remaining 58 OGs, we observed OG0006371 and OG0015023 containing genes from only 17 and 21 species, respectively. In OG0006371 we found Glyma.05G011200, an unannotated gene preferentially expressed in seed, flower, and nodule (Supplementary Figure 5). In OG0015023 we found Glyma.05G140300, a gene that encodes a small subunit of serine palmitoyltransferase-like (SPT-like) that is highly expressed in seeds. The SPT complex catalyzes the first and rate-limiting step in sphingolipid biosynthesis (M. Chen et al. 2006; P. Liu et al. 2023). In general, lipid metabolism genes related to adaptive responses including signaling and response biotic and abiotic stresses are found in OGs not shared by all species.

**Figure 6.**
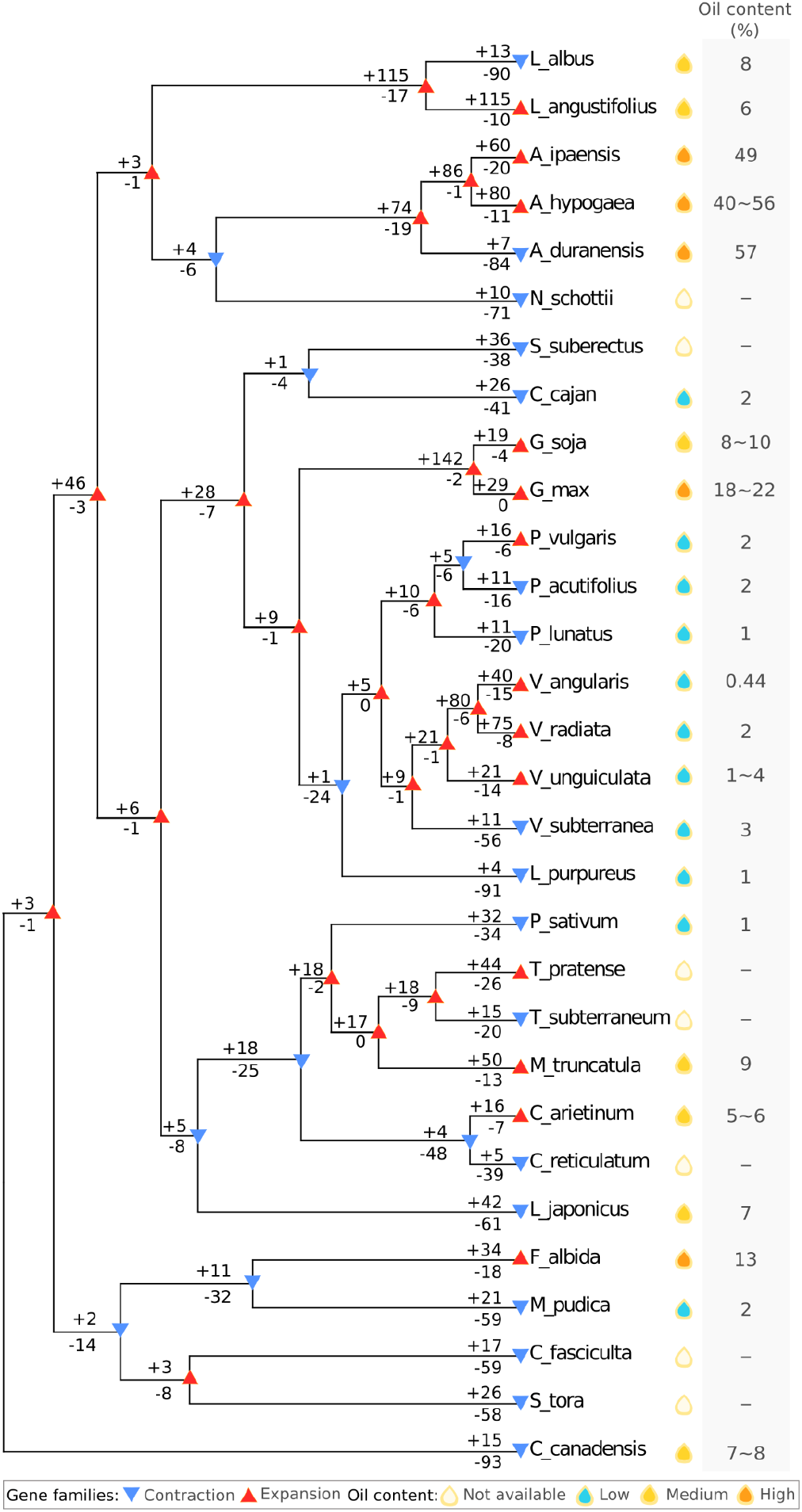
Species tree summarizing the number of orthogroups from gain families (∼8 TPM) with significant (p-value < 0.05) expansion or contraction across 30 legume species. Gain families are those with more genes in soybean than in common bean. Red and blue triangles refer to nodes/leaves with more expansions and contractions, respectively. The drops represent seed oil content in each species: Low: below 4.9%; Medium: between 5% and 10%; High: more than 10%. Oil content source: *A. duranensis* (L. Huang et al. 2012); *A. hypogaea* (Shasidhar et al. 2017); *A. ipaensis* (Grosso, Nepote, and Guzmán 2000); *C. cajan* (Sharma, Nidhi, and Preeti 2011); *C. canadensis* (Duke and Ayensu 1985); *C. arietinum* (Zia-Ul-Haq et al. 2007); *F. albida* (Hassan, Umar, and Yuguda 2007); *G.max* (Patil et al. 2018); *G. soja* (Patil et al. 2018); *L. purpureus* (Hossain et al. 2016); *L. japonicus* (Dam et al. 2009); *L. albus* (Bhardwaj, Hamama, and van Santen 2004); *L. angustifolius* (Lemus-Conejo et al. 2023); *M. truncatula* (Y. Song et al. 2017); *M. pudica* (Grygier et al. 2022); *P. acutifolius* (Bhardwaj, Hamama, and van Santen 2004); *P. lunatus* (Palupi et al. 2022); *P. vulgaris* (Sutivisedsak et al. 2011); *P. sativum* (Asen et al. 2023); *V. angularis* (Shweta and Katoch 2014); *V. radiata* (Zia-Ul-Haq, Ahmad, and Iqbal 2008); *V. subterranea* (Minka and Bruneteau 2000); *V. unguiculata* (Perchuk et al. 2020). The CAFE plot was generated with cafeplotter ( https://github.com/moshi4/CafePlotter).

## 3. Conclusion

The intricate history of plant WGD and the retention of multiple gene copies pose a considerable challenge in pinpointing the causative genes for specific traits. We employed a comprehensive approach for the investigation of gene families related to oil traits, using gene expression and co-expression, conservation and mutation rates. We identified soybean and common bean genes involved in TAG pathways, unveiled novel candidates and explored their expression and functional divergence in seeds, shedding light on their roles in lipid metabolism. Our findings do not only contribute to understanding the genetic mechanisms governing lipid metabolism, but also provide valuable leads for targeted genome editing for crop improvement and biotechnology (Figure 3, Supplementary Table 22).

## 4. Materials and methods

### 4.1 Selection of genes related to oil traits and enrichment analysis

We assembled a comprehensive dataset of soybean genes linked to oil traits from various sources: Aralip (McGlew et al. 2015), SoyCyc (v.9.0): diacylglycerol and triacylglycerol biosynthesis; and triacylglycerol degradation (Brown et al. 2021), and genes known for their involvement in lipid metabolism, obtained from the Mapman database, accessed via PLAZA Dicots 5.0 (Van Bel et al. 2022). These lists were supplemented with genes obtained through a systematic manual curation (Supplementary Table 1). This compilation resulted in a comprehensive collection of 2,176 soybean genes potentially associated with oil traits. The complete collection of homologous gene families were obtained from PLAZA 5.0, allowing the identification of 567 families containing potential oil genes. When considering the presence of homologs within these families, we expanded this set to 7,706 and 4,236 soybean and common bean candidate genes, respectively. Enrichment analyses for GO terms and conserved protein domains was performed in PhytoMine (https://phytozome.jgi.doe.gov/phytomine/begin.do), using Benjamini-Hochberg multiple testing correction (max p-value: 0.05) and the following background sets: genes in oil-candidate homologous families (Supplementary Table 4 and Supplementary Table 5); all genes expressed in seeds with ∼1 TPM (Supplementary Table 11); gain-family genes with at least 8 TPM (Supplementary Table 13); all soybean protein-coding genes (Supplementary Table 19 and Supplementary Table 21).

### 4.2 RNA-seq data and gene expression analysis

We conducted a meticulous selection of RNA-Seq samples from the Soybean Expression Atlas (https://soyatlas.venanciogroup.uenf.br/) (Almeida-Silva, Pedrosa-Silva, and Venancio 2023). To ensure the selection of relevant samples, a filtering process was carried out using the following criteria to exclude samples that originated from: 1) indeterminate plant parts (e.g. whole plant, seedling, and unknown); 2) transgenic or mutant plants; 3) cultivars other than Williams 82 and; 4) specific treatments (e.g. exposure to biotic and abiotic stresses). Exceptions for criteria 2 and 3 above include samples from mutants and varieties that exhibited specific advantages in the context of oil biosynthesis (e.g. Seed_jack_GmZF351, Seed_jack_GmZF352, seed_Thorne_wt_r5_r6 and seed_gmOleo1). This systematic curation resulted in a list of 605 samples (Supplementary table 23). Gene expression estimates were retrieved in TPM. We used the median TPM to investigate the gene expression patterns across the diverse array of samples. We also retrieved the tissue specificity index Tau (τ) (Kryuchkova-Mostacci and Robinson-Rechavi 2017; Yanai et al. 2005) available in the Soybean Expression Atlas (https://soyatlas.venanciogroup.uenf.br/). Expression data of *P. vulgaris* was obtained from public data PVGEA (O’Rourke et al. 2014). Expression levels were classified as low (TPM between 1 and 5), medium (TPM between 5 and 10), and high (TPM greater than 10).

### 4.3 Gene coexpression analysis

We used the R package BioNERO (Almeida-Silva and Venancio 2022) to construct a coexpression network of gain-family genes with at least 8 TPM in seeds. Data preprocessing included replacing missing values with 0, removal of non-expressed genes (median minimal expression = 1), removal outlying samples using the BioNero standard method (i.e. standardized connectivity - Z.K < 2), and adjusting for confounding artifacts to avoid spurious correlations. This process resulted in the exclusion of 35 samples. We used the WGCNA algorithm (Langfelder and Horvath 2008; B. Zhang and Horvath 2005) to compute the gene coexpression network. Hub genes were identified by combining two metrics: correlation of a gene to its module eigengene (module membership > 0.8) and sum of connection weights of a gene to all other genes in the module (degree; top 10% genes with highest degree). Network plots were generated using Cytoscape (Shannon et al. 2003).

### 4.4 Analysis of duplicated gene pairs

Protein sequences (.faa) and annotation data (.gff3) from Soybean (*G. max*, Wm82.a4.v1) and Common Bean (*P. vulgaris*, V2.1) were obtained from PLAZA 5.0 and Phytozome V12, respectively (Goodstein et al. 2012; Van Bel et al. 2022). Pairwise comparisons of soybean and common bean predicted proteins were conducted with Diamond 0.9.14 (Buchfink, Reuter, and Drost 2021). We used the DupGen Finder tool (Qiao et al. 2019) to classify gene duplication modes in one of five categories: dispersed (DD), proximal (PD), tandem (TD), transposed (TRD), and WGD (Supplementary Table 24). We computed nonsynonymous substitutions per nonsynonymous site (K_a_), synonymous substitutions per synonymous site (K_s_), and the K_a_/K_s_ ratio for all identified gene pairs with the calculate_Ka_Ks_pipeline script (Qiao et al. 2019). K_s_ peaks were predicted using the doubletrouble R package (Almeida-Silva and Van de Peer 2024).

### 4.5 Analysis of legume orthogroups

A diverse collection of proteome datasets spanning 30 distinct legume species gathered from Phytozome (V12/V13) (Goodstein et al. 2012), PLAZA 5.0 (Van Bel et al. 2022), GigaDB (http://gigadb.org/), Kazuza genome database (Shirasawa et al. 2014), DRYAD (Vision 2010) and Legumepedia (Garg et al. 2022). When multiple splicing isoforms were present for a gene, only the longest isoform was retained. We used OrthoFinder 2.5.2 (Emms and Kelly 2019) to infer OGs using the parameters -S diamond-ultra-sens (Buchfink, Reuter, and Drost 2021), -M msa, and -T iqtree (iqtree.org). Orthofinder results were explored using the cogeqc R/Bioconductor package (Almeida-Silva and Van de Peer 2023). We used CAFE5 (Mendes et al. 2020) to find expansions and contractions in the OGs (-k 3). As input, we used a time-calibrated ultrametric species tree (Kumar et al. 2022) and the gene counts from each species in each OG. The time-calibrated species tree was calculated using the script make_ultrametric.py based on the SpeciesTree-rooted from Orthofinder and root age (-r 68) available in TimeTree (Kumar et al. 2022). Large OGs indicated by CAFE5 were removed. The key parameters lambda (λ) and mi (μ) were estimated by running CAFE 50 times. The selection of optimal parameters was guided by the best maximum-likelihood estimate. CAFE results were visualized using the cafeplotter tool (https://github.com/moshi4/CafePlotter).

## 5. CRediT authorship contribution statement

Dayana K. Turquetti-Moraes: Conceptualization, Methodology, Validation, Formal analysis, Data curation, Investigation, Writing – original draft. Cláudio Benício Cardoso-Silva: selection of legume species and proteome data curation. Fabricio Almeida-Silva: Writing – review & editing, Methodology. Thiago M. Venancio: Conceptualization, Supervision, Project administration, Resources, Funding acquisition, Writing – original draft, Writing – review & editing.

## 6. Acknowledgements

This work was supported by Fundação Carlos Chagas Filho de Amparo à Pesquisa do Estado do Rio de Janeiro (FAPERJ), Coordenação de Aperfeiçoamento de Pessoal de Nível Superior—Brazil (CAPES, Finance Code 001) and Conselho Nacional de Desenvolvimento Científico e Tecnológico (CNPq). The funding agencies had no role in the design of the study and collection, analysis and interpretation of data and in writing.

## 7. Data and code availability

To ensure that all findings in this manuscript are reproducible, all data and code have been deposited on https://github.com/Dayana-Turquetti/Genes_related_to_oil_traits.

