## Supplementary figures for "Multiomic analysis of genes related to oil traits in legumes provide insights into lipid metabolism and oil richness in soybean"

[2] Laboratório Nacional de Ciência e Tecnologia do Bioetanol, Centro Nacional de Pesquisa em Energia e Materiais, Universidade de Campinas, São Paulo, SP, Brazil.

[3] Department of Plant Biotechnology and Bioinformatics, Ghent University, 9052 Ghent, Belgium.

[4] VIB Center for Plant Systems Biology, VIB, 9052 Ghent, Belgium.

* TMV: Laboratório de Química e Função de Proteínas e Peptídeos, Centro de Biociências e Biotecnologia, Universidade Estadual do Norte Fluminense Darcy Ribeiro. Av. Alberto Lamego 2000, P5, sala 217, Campos dos Goytacazes, RJ, Brazil.

### Supplementary Figures

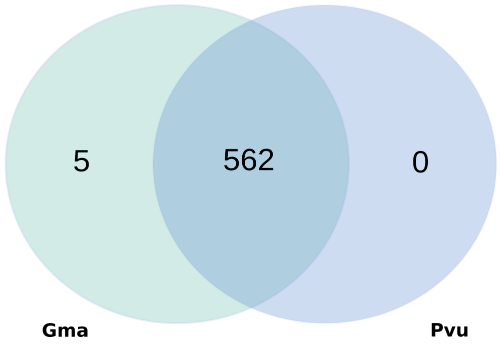


**Supplementary Figure 1**. Candidate oil homologous families for soybean (Gma) and common bean (Pvu). In total, we found 567 families of which 562 contain at least one common bean homolog, except HOM05D015518 (Glyma.13G001800; Glyma.20G068000), HOM05D006604 (Glyma.01G103450; Glyma.16G133700), HOM05D031525 (Glyma.07G184950; Glyma.08G064400), HOM05D130031 (Glyma.02G006100) and HOM05D039847 (Glyma.02G281400; Glyma.02G281500; Glyma.14G033100). Venn diagram was generated using Venn diagram ( https://bioinformatics.psb.ugent.be/webtools/Venn/).

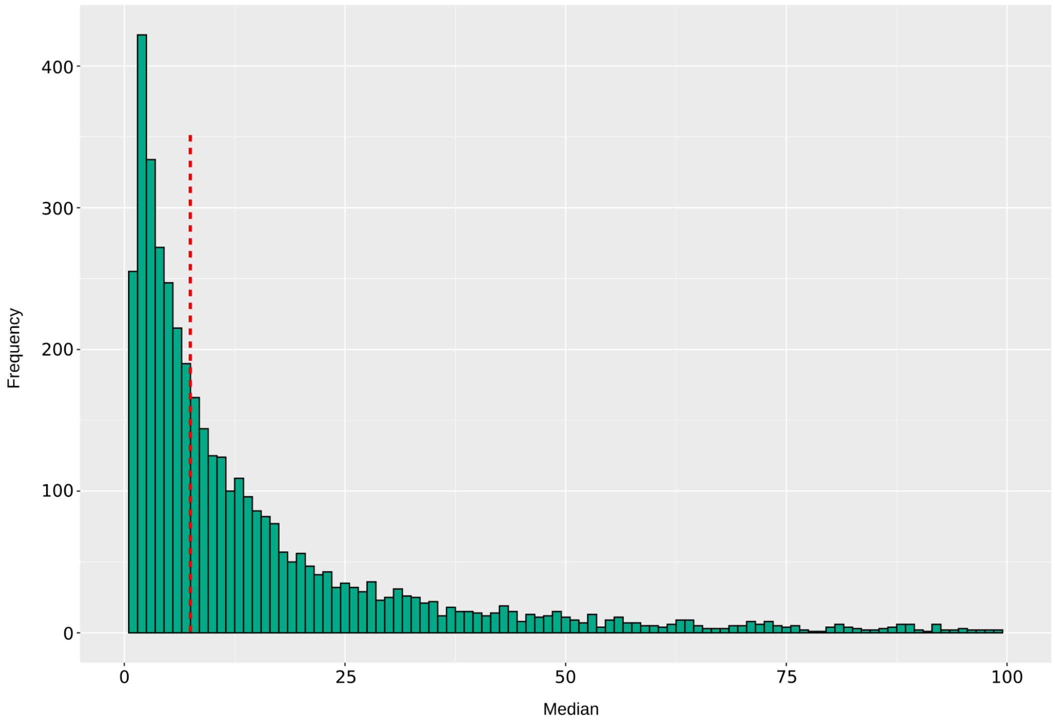


**Supplementary Figure 2**. Frequency distribution of maximum median TPM values for genes expressed in soybean seeds and seed subregions. A dashed red line indicates the threshold of 8 TPM.

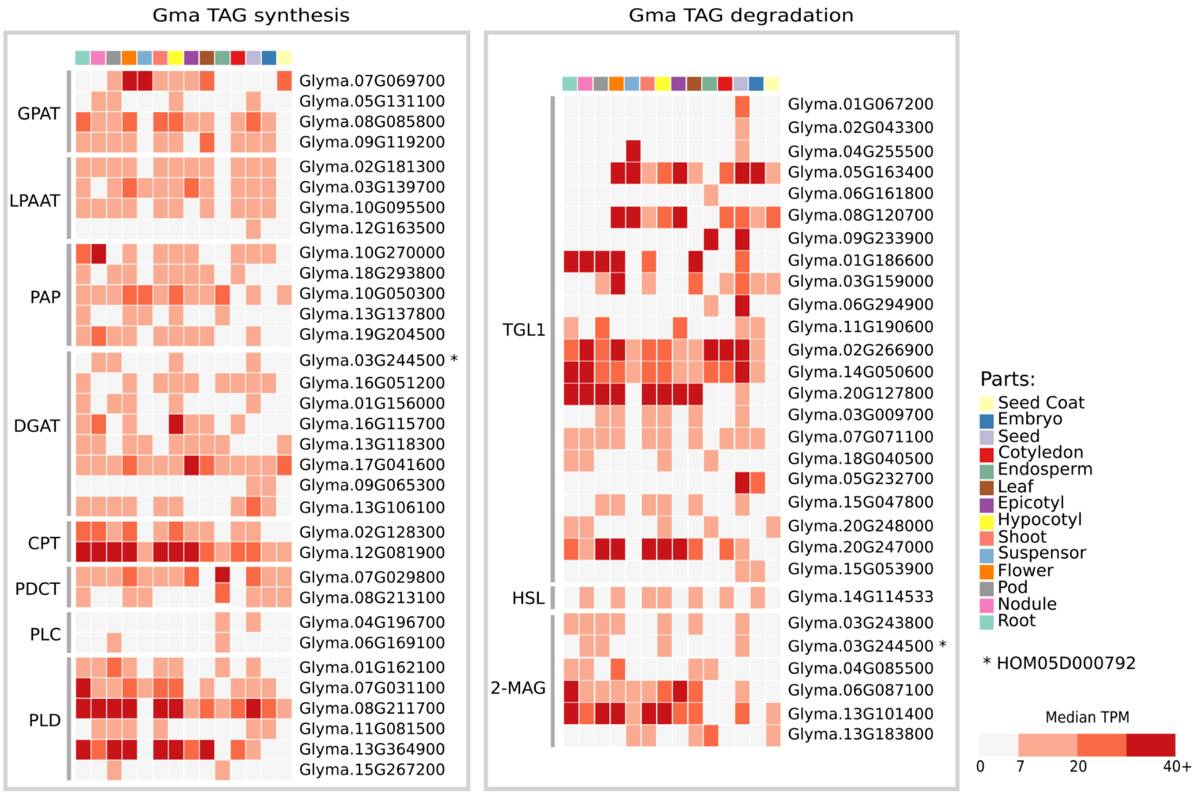


**Supplementary Figure 3**. Expression of soybean TAG pathway genes with at least 8 TPM in seeds or seed parts. **TPM:** transcripts per million. **GPAT**: acyl-CoA glycerol-3-phosphate acyltransferase; **LPAAT**: acyl-CoA lysophosphatidic acid acyltransferase; **PAP**: phosphatidic acid phosphatase; **DGAT**: acyl-CoA diacylglycerol acyltransferase; **PLC**: phospholipase C; **PLD**: phospholipase D; **CPT**: cytidine diphosphate-choline diacylglycerol cholinephosphotransferase; **PDCT**: phosphatidylcholine diacylglycerol cholinephosphotransferase; **TGL1**: triacylglycerol lipase; **HSL**: hormone-sensitive lipase; **2-MAG**: 2-monoacylglycerol acylhydrolase.

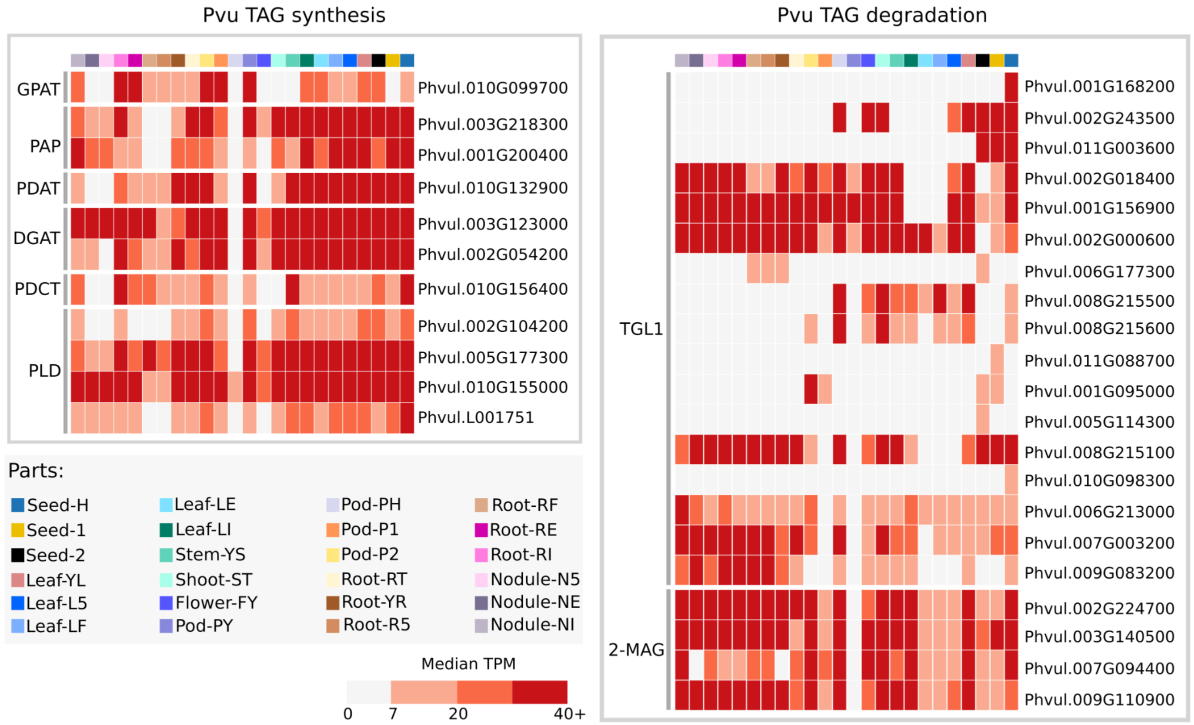


**Supplementary Figure 4**. Expression of common bean TAG pathway genes (at least 8 TPM in seeds). **TPM**: transcripts per million. **Seed-H**: seed heart stage; **Seed-1**: seeds with ~50mg; **Seed-2**: seeds with ~150mg; **Leaf-YL**: fully expanded second trifoliate leaf; **Leaf-L5**: leaf tissue collected 5 days after plants were inoculated with effective rhizobium; **Leaf-LF**: leaf tissue from fertilized plants; **Leaf-LE**: leaf tissue collected 21 days after plants were inoculated with effective rhizobium; **Leaf-LI**: leaf tissue collected 21 days after plants were inoculated with ineffective rhizobium; **Stem-YS**: all stem internodes above the cotyledon collected at the second trifoliate stage; **Shoot-ST**: shoot tip; **Flower-FY**: young flowers; **Pod-PY**: young pods; **Pod-PH**: pods 9cm long; **Pod-P1**: pods between 10 and 11 cm long; **Pod-P2**: pods between 12 and 13 cm long; **Root-RT**: root tips; **Root-YR:** whole roots at the second trifoliate stage of development; **Root-RS**: whole roots separated from 5 days old pre-fixing nodules; **Root-RF**: whole roots from fertilized plants; **Root-RE:** whole roots separated from fix+ nodules collected 21 days after inoculation; **Root-RI**: whole roots separated from fix- nodules collected 21 days after inoculation; **Nodule-NS**: pre-fixing effective nodules collected 5 days after inoculation; **Nodule-NE**: effectively fixing nodules collected 21 days after inoculation; **Nodule-NI**: ineffectively fixing nodules collected 21 days after inoculation.

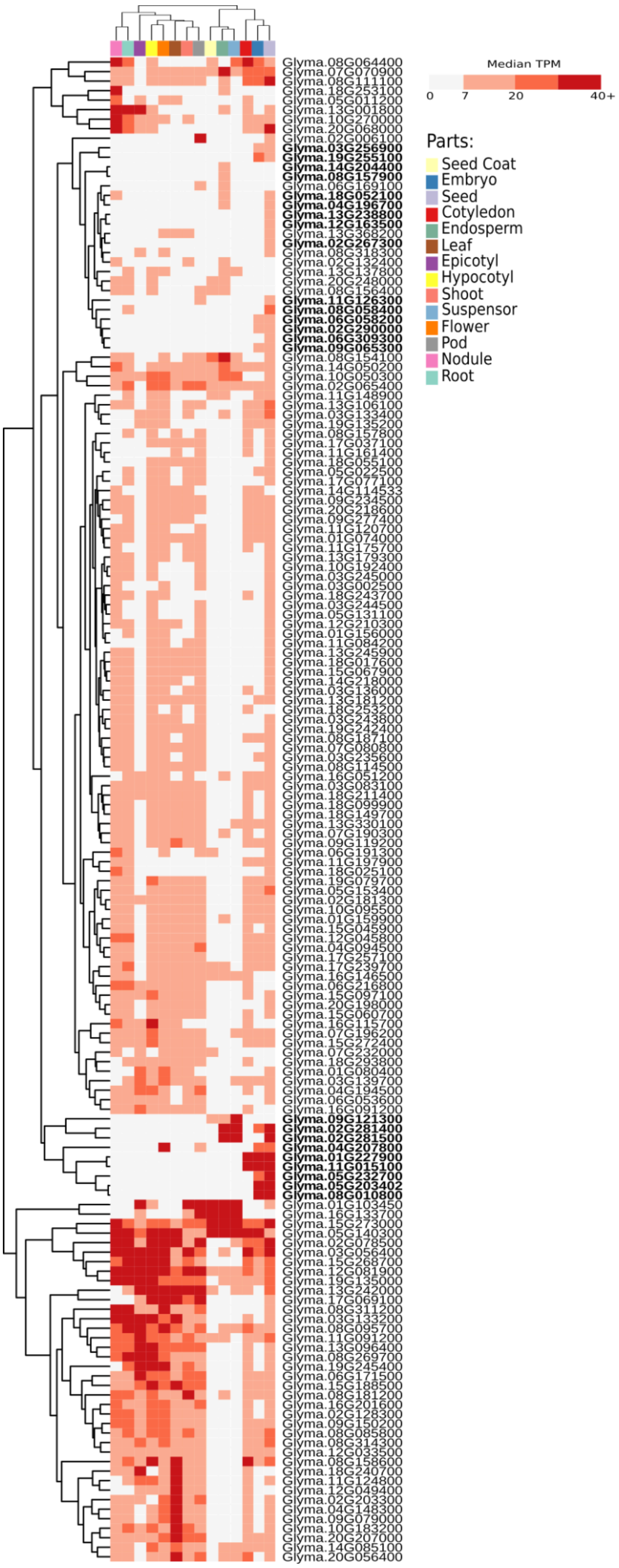


**Supplementary Figure 5**. Genes from 90 homologous families with expression in soybean but not in common bean seeds. Genes more expressed in seed or seed subregions than in other parts are marked in bold.

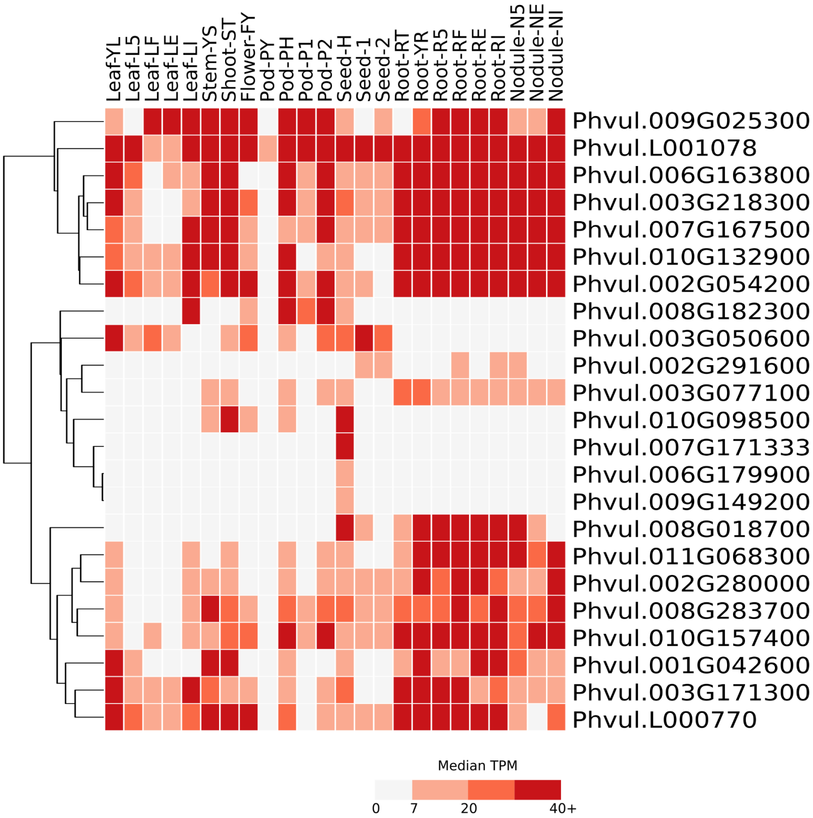


**Supplementary Figure 6**. Expression of common bean genes from the third category of genes with high expression in seeds. **TPM**: transcripts per million. **Seed-H**: seed heart stage; **Seed-1**: seeds with ~50mg; **Seed-2**: seeds with ~150mg; **Leaf-YL**: fully expanded second trifoliate leaf; **Leaf-L5**: leaf tissue collected 5 days after plants were inoculated with effective rhizobium; **Leaf-LF**: leaf tissue from fertilized plants; **Leaf-LE**: leaf tissue collected 21 days after plants were inoculated with effective rhizobium; **Leaf-LI**: leaf tissue collected 21 days after plants were inoculated with ineffective rhizobium; **Stem-YS**: all stem internodes above the cotyledon collected at the second trifoliate stage; **Shoot-ST**: shoot tip; **Flower-FY**: young flowers; **Pod-PY**: young pods; **Pod-PH**: pods 9cm long; **Pod-P1**: pods between 10 and 11 cm long; **Pod-P2**: pods between 12 and 13 cm long; **Root-RT**: root tips; **Root-YR:** whole roots at the second trifoliate stage of development; **Root-RS**: whole roots separated from 5 days old pre-fixing nodules; **Root-RF**: whole roots from fertilized plants; **Root-RE:** whole roots separated from fix+ nodules collected 21 days after inoculation; **Root-RI**: whole roots separated from fix- nodules collected 21 days after inoculation; **Nodule-NS**: pre-fixing effective nodules collected 5 days after inoculation; **Nodule-NE**: effectively fixing nodules collected 21 days after inoculation; **Nodule-NI**: ineffectively fixing nodules collected 21 days after inoculation.

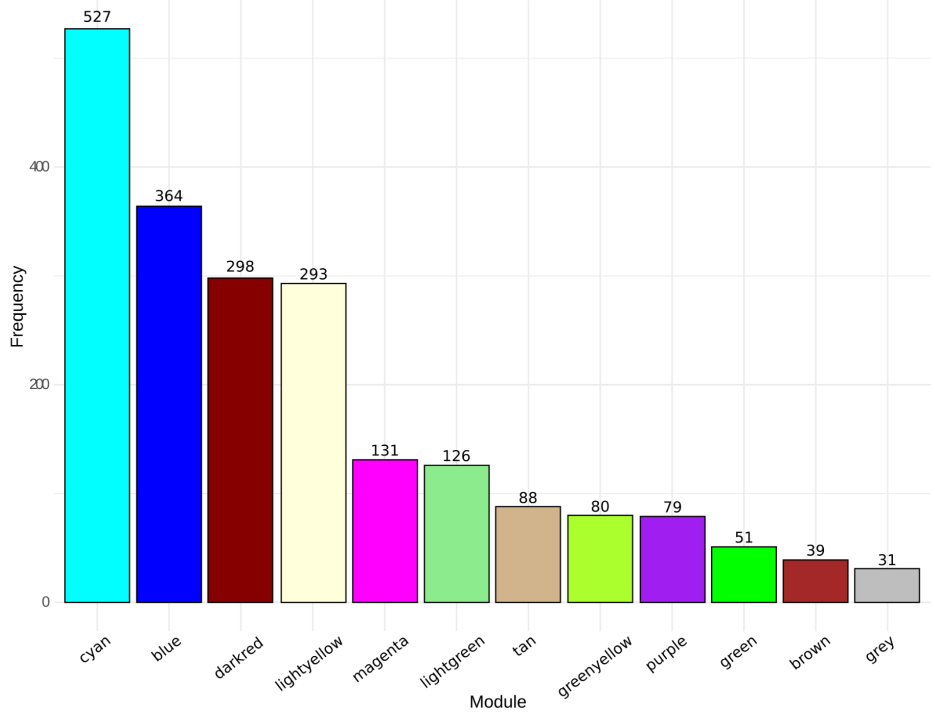


**Supplementary Figure 7**. Frequency of soybean oil genes in twelve co-expression modules.

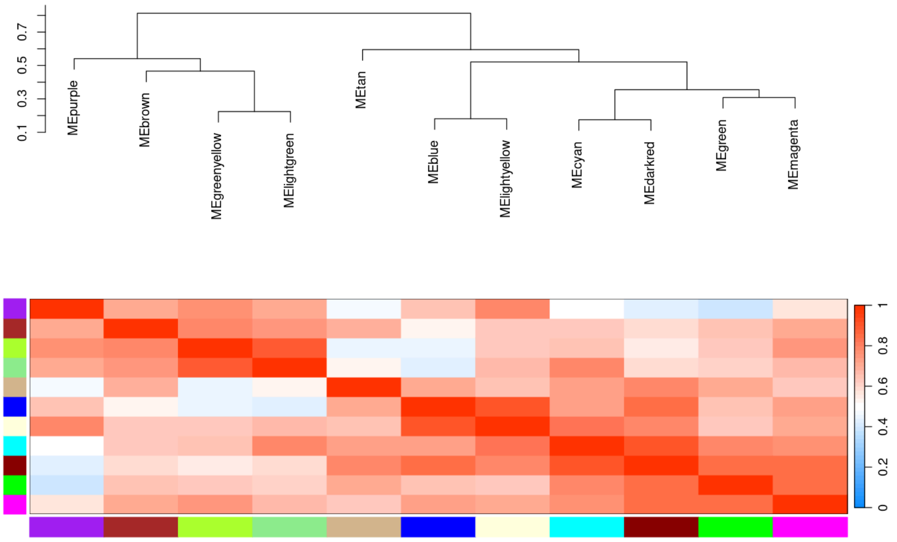


**Supplementary Figure 8**. Modules eigengene indicating the proximity (clusters) of the expression profile among eleven modules. Module eigengene is the first principal component of a principal component analysis, which summarizes the expression of the module.

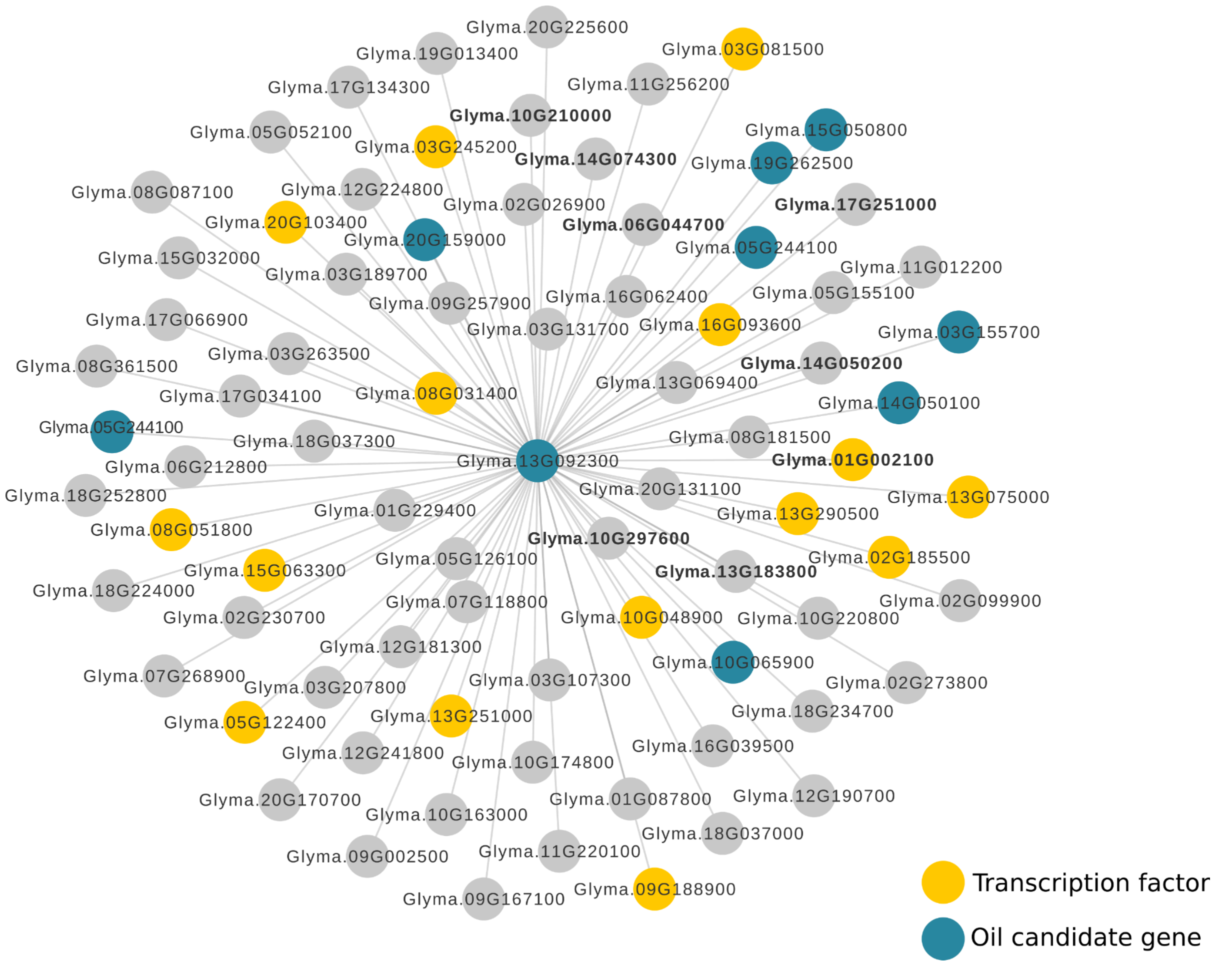


**Supplementary Figure 9**. Connections of the hub Glyma.13G092300. Transcription factors and previously reported oil candidate genes (Niu et al., 2020) are highlighted in yellow and green, respectively. The connections of these three hubs with relevant genes to lipid metabolism and TFs support their importance in oil-related traits. Genes reported as associated with lipid metabolism in Aralip or Mapman are marked in bold. Network was generated using Cytoscape (Shannon et al., 2003, available at https://doi.org/10.1101/gr.1239303).

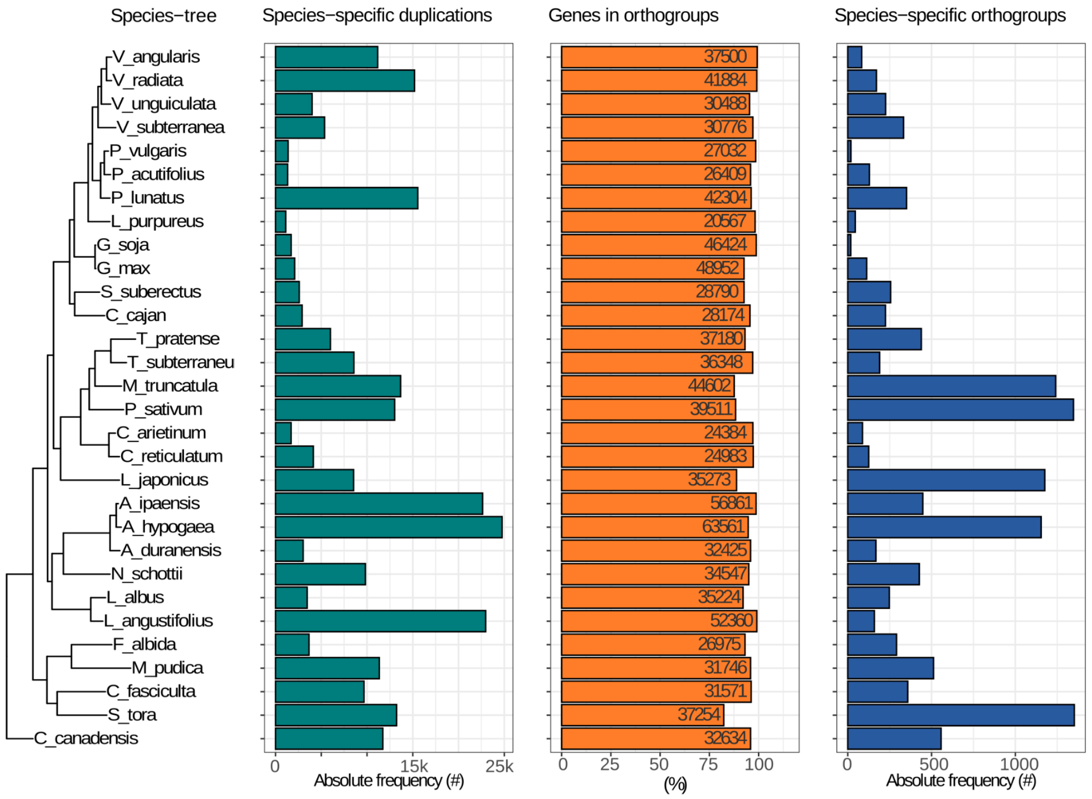


**Supplementary Figure 10.** Summary of the orthogroup analysis. From left to right, the panels contain the species-tree, absolute frequency of the species-specific duplications, percentage of the genes in orthogroups and absolute frequency of the species-specific orthogroups. The image was generated using Cogeqc R/Bioconductor package (Almeida-Silva and Van de Peer, 2023).
